# Test-time augmentation for deep learning-based cell segmentation on microscopy images

**DOI:** 10.1101/814962

**Authors:** Nikita Moshkov, Botond Mathe, Attila Kertesz-Farkas, Reka Hollandi, Peter Horvath

## Abstract

Recent advancements in deep learning have revolutionized the way microscopy images of cells are processed. Deep learning network architectures have a large number of parameters, thus, in order to reach high accuracy, they require massive amount of annotated data. A common way of improving accuracy builds on the artificial increase of the training set by using different augmentation techniques. A less common way relies on test-time augmentation (TTA) which yields transformed versions of the image for prediction and the results are merged. In this paper we describe incorporating the test-time argumentation prediction method into two major segmentation approaches used in the single-cell analysis of microscopy images, namely semantic segmentation using U-Net and instance segmentation using Mask R-CNN models. Our findings show that even using only simple test-time augmentations, such as rotation or flipping and proper merging methods, will result in significant improvement of prediction accuracy. We utilized images of tissue and cell cultures from the Data Science Bowl (DSB) 2018 nuclei segmentation competition and other sources. Additionally, boosting the highest-scoring method of the DSB with TTA, we could further improve and our method has reached an ever-best score at the DSB.

## Introduction

Identifying objects at the single-cell level is the starting point of most microscopy-based quantitative cellular image analysis tasks. Precise segmentation of the cell’s nucleus is a major challenge here. Numerous approaches have been developed, such as methods using mathematical morphology^1^ or differential geometry^2,3^. More recently deep learning has yielded a never-seen improvement of accuracy and robustness^4, 5, 6.^. Remarkably, Kaggle’s Data Science Bowl 2018 (DSB)^7^ was dedicated to nuclei segmentation and gave a great momentum to this field. Deep learning-based approaches have proved their effectiveness: practically all the teams used some type of a deep architecture in the first few hundred leaderboard positions. The most popular architectures included U-Net^4^, originally designed for medical image segmentation, and Mask R-CNN^8^, used for instance segmentation of natural objects.

Deep learning approaches for object segmentation require a large and often pixel-wise annotated dataset for training. This task relies on high-quality samples and domain experts to accurately annotate images. Besides, analysing biological images is challenging because of their heterogeneity and, sometimes, poorer quality compared to natural images. In addition, ground truth masks might be imperfect due to the annotator-related bias, which introduces further uncertainty. Consequently, a plethora of annotated samples is required, making object segmentation a laborious process. One of the techniques utilized to improve the model is data augmentation^9^ of the training set. Conventionally, a transformation (i.e. rotation, flipping, noise addition etc.) or a series of transformations are applied on the original images. Data augmentation has become the *de facto* technique in deep learning, especially in the case of heterogeneous or small datasets to improve the accuracy of cell-based analysis.

To improve performance, another possibility relies on augmenting not only the training dataset, but also the test dataset, thus performing the prediction on the original, as well as on the augmented versions of the image, and merging the predictions; this approach is called ***test-time augmentation*** (Figure 1). This technique was successfully used in image classification tasks^10^, for aleatoric uncertainty estimation^11^ and MRI slices/MRI volumes segmentation^12^. A theoretical formulation^12^ of test-time augmentation has been described recently, their experiments show that TTA helps to get rid of overconfident incorrect predictions. Additionally, a framework^13^ for quantifying the uncertainty of the DNN model for diagnosing diabetic retinopathy based on test-time data augmentation was proposed. Its disadvantage is the increased prediction time, as it is run not only on the original image, but on all its augmentations as well.

**Figure 1.**
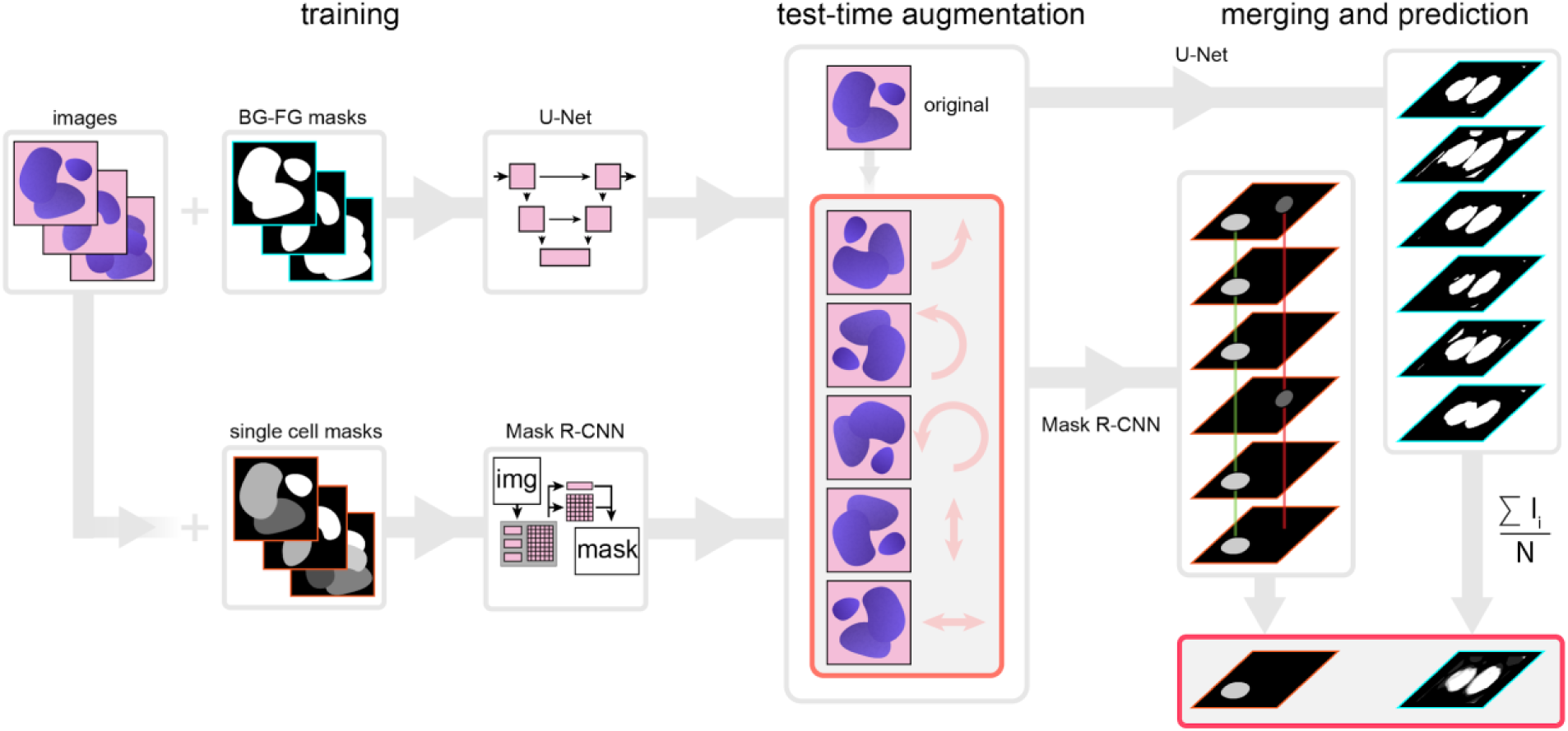
Principle of the proposed test-time augmentation techniques. Several augmented instances of the same test images were predicted and the results were transformed back and merged. In the case of U-Net, pixel-wise majority voting was applied, while for Mask R-CNN, a combination of object matching and majority voting was used.

In the current paper we assess the impact and describe cases of utilizing test-time augmentation for deep-learning models trained on microscopy datasets. We have trained deep learning models for both semantic segmentation (when the network only distinguishes the foreground from the background, using the U-Net architecture) and instance segmentation (when the network assigns labels to separate objects, using the Mask R-CNN architecture) (Figure 1). In conclusion, test-time augmentation has outperformed single instance predictions at each test cases, and could further improve the current best result of the DSB, as demonstrated by the improved score, changing from 0.633 to 0.644.

**Figure 2.**
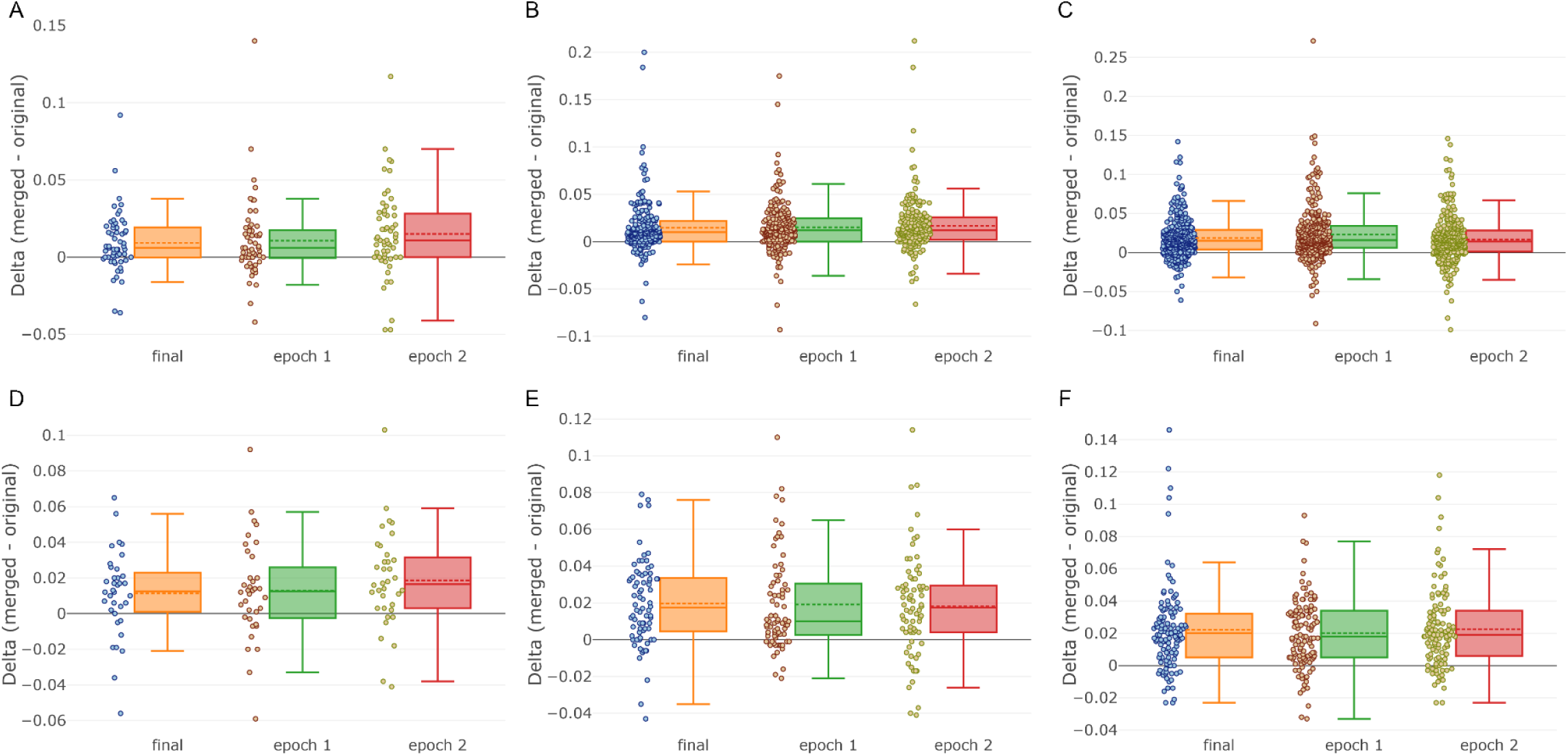
TTA performance for Mask R-CNN. TTA performance (*delta* = *merged* − *original*). Each point represents an image. Dashed line – mean, solid line – median. A | Fluorescent set 5. B | Fluorescent set 15. C | Fluorescent set 30. D | Tissue set 5. E | Tissue set 15. F | Tissue set 30.

**Figure 3.**
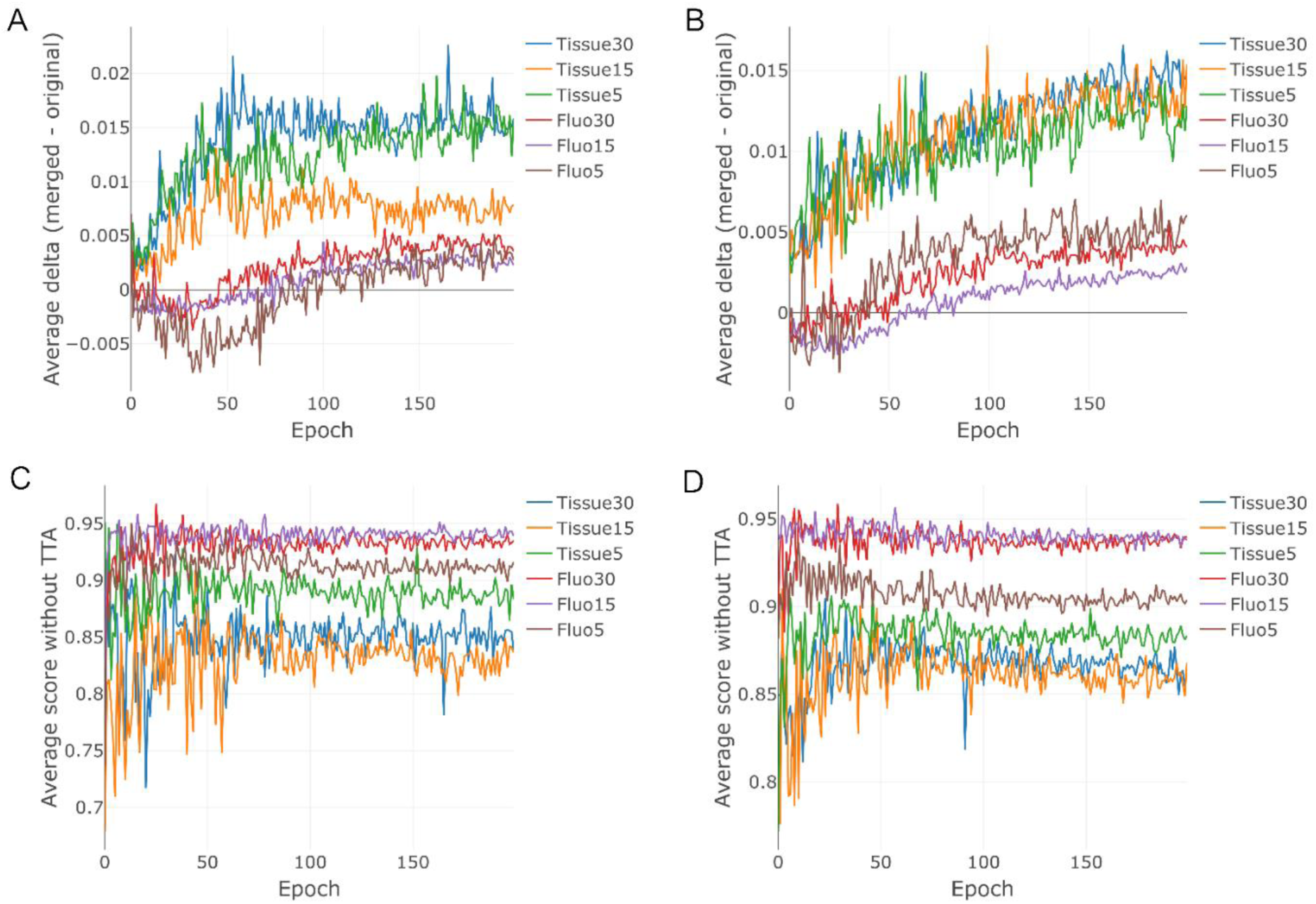
Average performance for U-Net with different training and test augmentations. A | Average TTA performance trained without augmentations over epochs. B | Average TTA performance trained with augmentations over epochs. C | Average performance without TTA without augmentations during training. D | Average performance without TTA with augmentations during training.

## Methods

### Dataset acquisition and description

We have collected two datasets: fluorescent microscopy images (further referred to as ‘fluorescent’ dataset) and histopathology images (further referred to as ‘tissue’ dataset). Most of the images have come from the stage 1 train/test data of Data Science Bowl 2018. We also used additional sources^14,15,16,17,18,19,20^ and other data published in the discussion thread ‘Official External Data Thread’ (https://www.kaggle.com/c/data-science-bowl-2018/discussion/47572) related to DSB 2018. The images were labelled by experts using the annotation plugins of ImageJ/Fiji and Gimp. Both datasets were divided into three holdout train\test sets: approximately 5%, 15%, 30% (further referred to as ‘5’, ‘15’ and ‘30’ in the dataset name, respectively) of uncropped images were held out as the test set. The test sets did not intersect.

We used the same augmentations (horizontal and vertical flip, 90°, 180° and 270° rotations) for training both architectures. The images were cropped to the size of 512×512 pixels. Crops from the same image were used only in either the train or test set. Images with a resolution lower than 512×512 were resized to that particular size. Sample images are shown in Figure 4.

**Figure 4.**
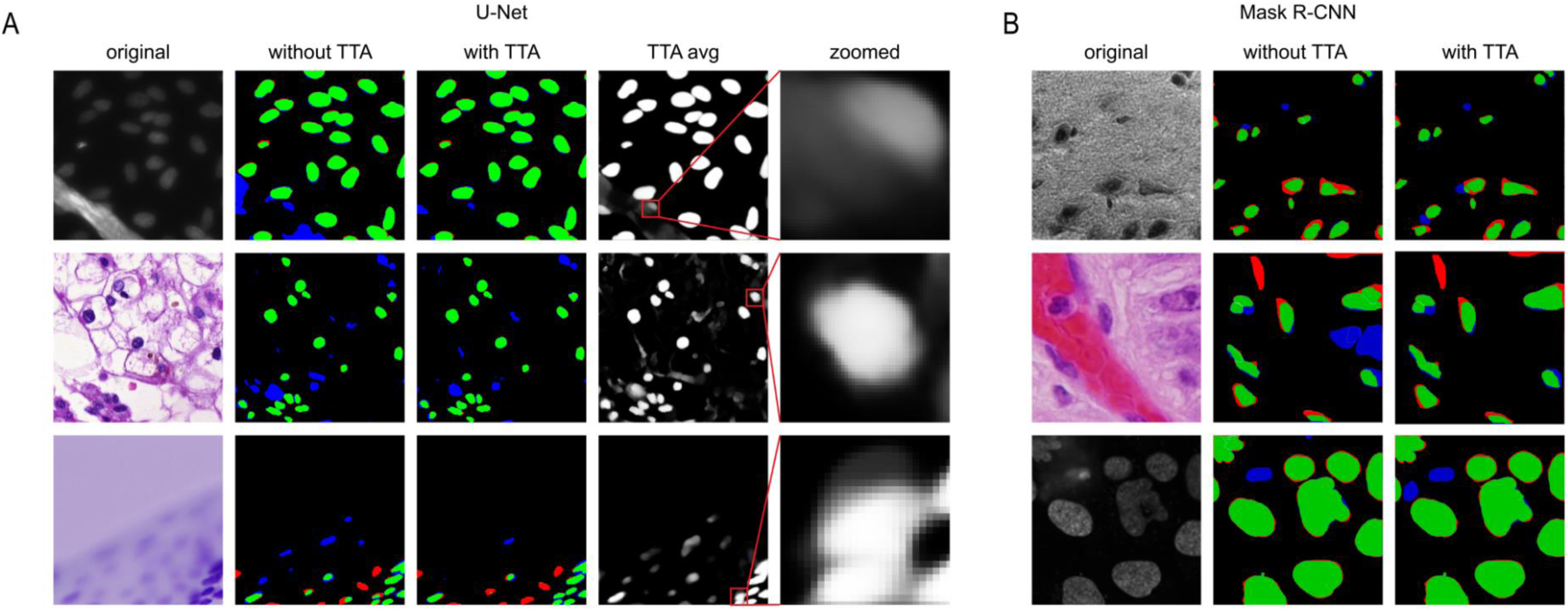
Examples of predictions. A | U-Net predictions. First column – original image, second column – predictions without TTA compared to ground truth, third column – predictions with TTA compared to ground truth. Red marks indicate false negative, green marks indicate true positive and blue marks indicate false positive. Fourth column – averaged TTA predictions before thresholding, fifth column – zoomed insets from the previous column. B | Mask R-CNN predictions. Columns are as the first three columns in A.

### Deep learning models and training

These augmented and cropped training data were used to train the models. For each dataset (5, 15 and 30 holdouts for both fluorescent and tissue images) separate models were trained. Additionally, we also trained U-Net without augmented data to analyse TTA performance on such a network as well.

Mask R-CNN (implementation^21^) is an extension of Faster R-CNN, the architecture for object detection. Solutions based on Mask R-CNN outperform the COCO 2016 challenge winners and finished at the third place in Kaggle Data Science Bowl 2018^7^. The architecture of Mask R-CNN incorporates the following main stages: (1) Region proposal network (RPN) to propose candidate bounding boxes. It uses a backbone: a convolutional neural network which serves as a feature extractor. In this implementation it is possible to use *resnet50* or *resnet101* as a backbone, we used *resnet101*. (2) Network head layers: they predict the class, box offset and an output binary mask for each region of interest (RoI). Masks are generated for each class without competition between classes.

The network, following the strategy described by Hollandi *et al.5*, was trained for 3 epochs for different layer groups: first, all network layers were trained at a learning rate of 10^−3^, then training was restricted to ResNet stage 5 (ResNet consists of 5 stages, each with convolution and identity blocks including 3 convolutional layers per block) and head layers at a learning rate of 5 × 10^−4^, and finally only the head layers were trained at a learning rate of 10^−4^. The model was initialized with pre-trained weights (https://github.com/matterport/Mask_RCNN/releases/download/v1.0/mask_rcnn_coco.h5) on the COCO dataset. The loss function of the architecture was binary cross-entropy with ADAM^22^ (Adaptive Moment Estimation) solver, batch size 1, the number of iterations being equal to the train set size.

U-Net (implementation^23^) is an architecture originally designed to process biological images, which proved to be efficient, even when utilizing small training datasets. U-Net based solutions won the 2015 ISBI cell tracking challenge^4^ and Kaggle Data Science Bowl 2018. Its architecture consists of two main parts: (1) a down-sampling convolution network or encoder by which we obtain the feature representation of the input image, and (2) an up-sampling convolution network or decoder, which produces the segmentation from a feature representation of the input image.

We trained U-Net for 200 epochs at a constant learning rate of 3 × 10 ^−4^, and used a binary cross-entropy loss function with ADAM solver, batch size 1, the number of iterations being equal to the train set size.

Both U-Net and Mask R-CNN implementations are based on the deep learning framework Keras with Tensorflow backend. The training computations were conducted on a PC with NVIDIA Titan Xp GPU, 32 GB RAM and Core-i7 CPU.

### Test-time augmentation

Test-time augmentation includes four procedures: augmentation, prediction, dis-augmentation and merging. We first apply augmentations on the test image. These are the same as the augmentations previously applied on the training dataset. We predict on both the original and the augmented images, then we revert the transformation on the obtained predictions; this process is referred to as dis-augmentation. For example, when the prediction was performed on a flipped or rotated image, we restore the obtained prediction to its original orientation. The final merging step is not straightforward in case of Mask R-CNN as the architecture is instance aware, thus the merging method has to handle instances. We have developed an extended merging method inspired by one of DSB 2018 solutions^24^ (Figure 1, right). For each detected object from the original image, we find the same detected object in the augmented images by calculating intersection over union (IoU) between the masks. The minimum IoU threshold used to decide whether the objects found are the same is 0.5. We iterate over all detected objects to find the best match. An object should be present in the majority of the images to be included as a final mask. Next, we check the first augmented image for any remaining unused objects (a possible scenario when an object is not detected in the original image but is detected in any of the augmented ones), and look for matching unassigned objects on other augmentations. Next, we check the second augmented image for detected objects and perform the same operations. We repeat this process until the majority voting criterion can be theoretically satisfied (in half of the images at a maximum). An average binary object mask is created by majority pixel voting on paired objects.

For U-Net the merging process is straightforward as it is not instance aware, so we simply sum and average all the dis-augmented probability maps. This results in a floating point image that needs to be converted to a binary mask. A simple element-wise thresholding at the value of 0.5 converts the soft masks binary (Figure 1., right).

### Test-time augmentation evaluation

We have evaluated the test-time augmentation model on our test dataset predictions (see the previous section for details) compared to ground truth masks using the following evaluation strategies.

In case of Mask R-CNN we used the same metric as at the Data Science Bowl 2018. It calculates the mean average precision (mAP) at different intersection over union (IoU) thresholds. The thresholds (t) are in the range of [0.5, 0.95] with a step of 0.05. An object is considered true positive when the IoU with ground truth is greater than the threshold, false positive when the predicted object has no associated ground truth object or the overlap is smaller than the threshold and false negative when the ground truth object has no associated predicted object.

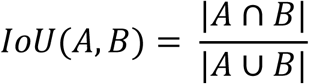

Thus, mAP for an image is calculated as follows:

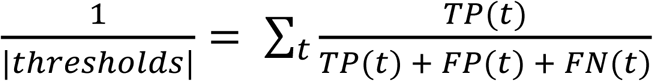

Next, we calculate the average for all images in the test set. The final score is a value between 0 and 1.

U-Net predictions were evaluated using the intersection over union metric, executed at the pixel level. We summed up the prediction and ground truth binary masks then we simply counted the pixels that are greater than one (that is the intersection) and divided the resulting values with the number of pixels greater than zero. The resulting value is a score ranging from 0 to 1.

As described above, we have evaluated the predictions with applying TTA (*merged*) and without applying TTA (*original*). Next, we have evaluated TTA’s performance by calculating the difference *delta* = *merged* − *original*.

## Results

In the case of Mask R-CNN, TTA on average has provided an improved performance for all dataset splits and for all model checkpoints. The average mAP score *delta* is about 0.01 for all “Fluorescent” and “Tissue_5” sets and 0.02 for the other sets. In all scenarios, TTA has positively affected the score for most of the images (Figure 2). In the case of U-Net, we have evaluated the performance at each epoch during training. For the “Tissue” datasets TTA demonstrated a performance gain for all epochs. In case of the “Fluorescent” datasets, a slight decline in the performance of TTA was observed during early (first 30-50) epochs which has turned positive after further training (Figure 3, A and B). After about epoch 50, the performance without TTA was seen to fluctuate without a clear trend in all cases (Figure 3, C and D), while the performance with TTA tended to rise for almost all cases, except in the case of the “Tissue” dataset, where no augmentations were used for training (Figure 3 A). For some images TTA has changed the final prediction result in a significantly positive manner. Examples of such cases for both U-Net and Mask R-CNN are shown in Figure 4.

Applying TTA on the DSB2018 (stage2) test set of images, it was found to further improve performance significantly, surpassing the best performing method^5^ by 0.011 (nearly 2%) in the DSB scoring metric which is identical to the mAP used in this paper and the output of which was instance segmented masks (Figure 5). Results without TTA and delta values for each set can be found in Supplementary (Supplementary Table 1. – U-Net when augmentations during training were used, Supplementary Table 2. – U-Net when augmentations during training were not used and Supplementary Table 3. – for Mask R-CNN).

**Figure 5.**
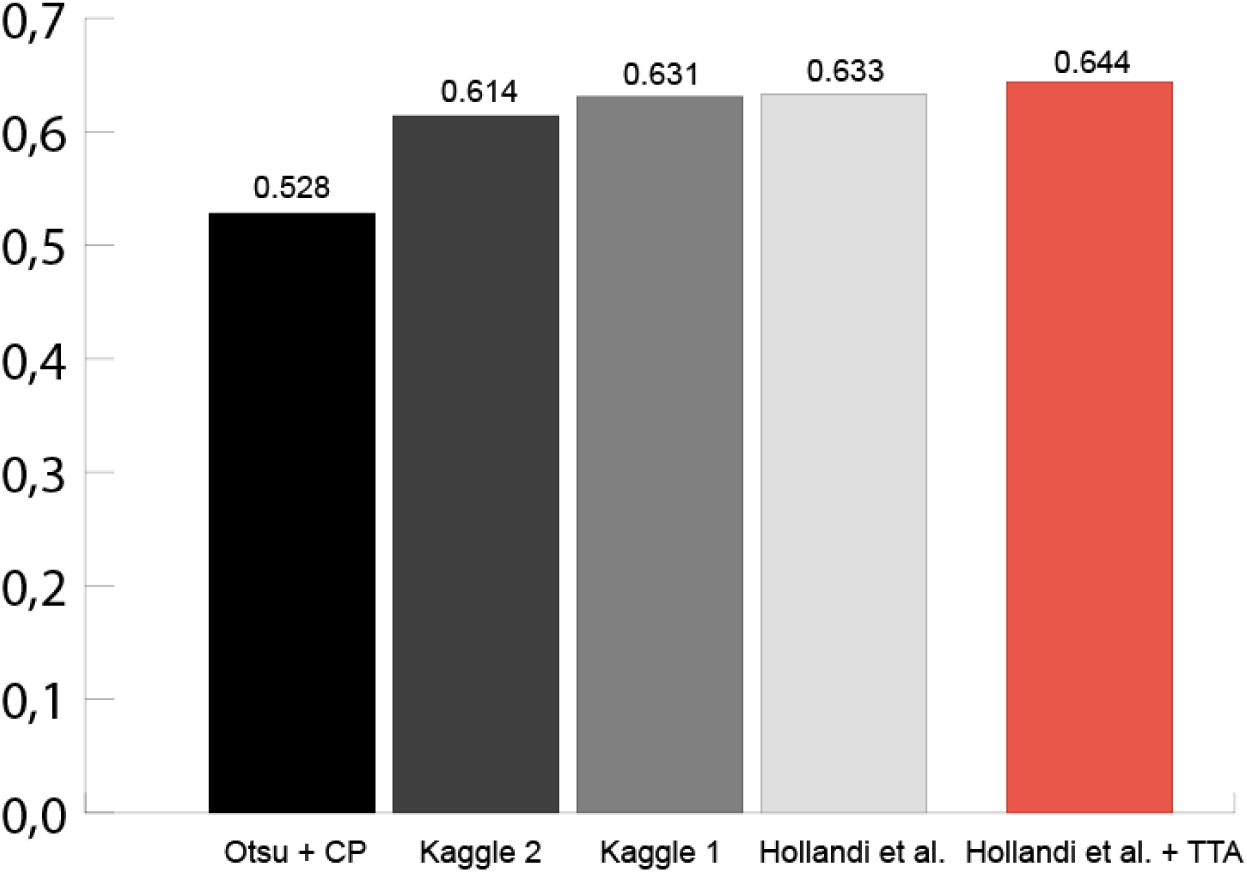
DSB Stage 2 scores for various methods (CellProfiler, Kaggle DSB 2018 2nd and 1st places, Hollandi et al.^5^ method and the same method with TTA). The red bar shows the highest score.

## Conclusions

We have performed experiments to estimate test-time augmentation’s performance for two popular deep learning frameworks trained to segment nuclei in microscopy images. Our results indicate that on average TTA can provide higher segmentation accuracy compared to only predicting on original images, even though for some images the results might be marginally worse.

TTA mostly affects the objects’ borders but in uncertain cases it can help to fit whole objects (remove false positives or add true positives, especially in case of Mask R-CNN). In the case of U-Net, TTA has rarely had a significant effect on segmentation results. Overall, in most cases, TTA improves segmentation accuracy. The main use case of TTA is examination of uncertain regions in segmentation. However, the high cost of TTA, related to the fact that multiple times more predictions are required for the same object, is also an issue to be considered. Therefore, TTA is mainly recommended for use when the basic cost of prediction is low.

## Authors’ contributions

N.M., B.M., R.H., A. K.-F. and P.H. wrote the manuscript. N.M, B.M and R.H. performed the experiments. R.H. collected the dataset. R.H. and N.M. prepared the figures for the paper.

## Acknowledgements

NM, RH, BM, and PH acknowledge support from the LENDULET-BIOMAG Grant (2018-342) and from the European Regional Development Funds (GINOP-2.3.2-15-2016-00006, GINOP-2.3.2-15-2016-00037), N.M. is supported by the Doctoral School of Interdisciplinary Medicine, the University of Szeged and the Doctoral School of Computer Science at the National Research University Higher School of Economics.

The authors acknowledge an NVIDIA grant for TitanXp GPUs.

The authors thank Dora Bokor, PharmD, for proofreading the manuscript.

## References

1. Carpenter, A. E. et al. CellProfiler: image analysis software for identifying and quantifying cell phenotypes. Genome Biol. 7, R100 (2006).

2. Molnar, C. et al. Accurate Morphology Preserving Segmentation of Overlapping Cells based on Active Contours. Sci. Rep. 6, 32412 (2016).

3. Molnar, J., Molnar, C. & Horvath, P. An Object Splitting Model Using Higher-Order Active Contours for Single-Cell Segmentation. Advances in Visual Computing 24–34 (2016). doi: 10.1007/978-3-319-50835-1_3

4. Ronneberger, O., Fischer, P. & Brox, T. U-Net: Convolutional Networks for Biomedical Image Segmentation. Lecture Notes in Computer Science 234–241 (2015). doi: 10.1007/978-3-319-24574-4_28

5. Hollandi, R. et al. A deep learning framework for nucleus segmentation using image style transfer. bioRxiv 580605 (2019). doi: 10.1101/580605

6. Dobos, O., Horvath, P., Nagy, F., Danka, T. & Viczián, A. A deep learning-based approach for high-throughput hypocotyl phenotyping. bioRxiv 651729 (2019). doi: 10.1101/651729

7. Juan C. Caicedo, Allen Goodman, Kyle W. Karhohs, Beth A. Cimini, Jeanelle Ackerman, Marzieh Haghighi, CherKeng Heng, Tim Becker, Minh Doan, Claire McQuin, Mohammad Rohban, Shantanu Singh, Anne E. Carpenter. Nucleus segmentation across imaging experiments: the 2018 Data Science Bowl. Nature Methods (2019). doi: 10.1038/s41592-019-0612-7

8. He, K., Gkioxari, G., Dollar, P. & Girshick, R. Mask R-CNN. IEEE Trans. Pattern Anal. Mach. Intell. (2018). doi: 10.1109/TPAMI.2018.2844175

9. Krizhevsky, A., Sutskever, I. & Hinton, G. E. ImageNet classification with deep convolutional neural networks. Communications of the ACM 60, 84–90 (2017).

10. Matsunaga, K., Hamada, A., Minagawa, A. & Koga, H. Image Classification of Melanoma, Nevus and Seborrheic Keratosis by Deep Neural Network Ensemble. (2017).

11. Ayhan, M. S. & Berens, P. Test-time Data Augmentation for Estimation of Heteroscedastic Aleatoric Uncertainty in Deep Neural Networks. (2018).

12. Wang, G. et al. Aleatoric uncertainty estimation with test-time augmentation for medical image segmentation with convolutional neural networks. Neurocomputing 338, 34–45 (2019).

13. Ayhan, M. S. et al. Expert-validated estimation of diagnostic uncertainty for deep neural networks in diabetic retinopathy detection. medRxiv 19002154 (2019).

14. Brasko, C. et al. Intelligent image-based in situ single-cell isolation. Nature Communications 9, (2018).

15. Caicedo, J. C. et al. Evaluation of Deep Learning Strategies for Nucleus Segmentation in Fluorescence Images. doi: 10.1101/335216

16. Caie, P. D. et al. High-content phenotypic profiling of drug response signatures across distinct cancer cells. Mol. Cancer Ther. 9, 1913–1926 (2010).

17. Coelho, L. P., Shariff, A. & Murphy, R. F. Nuclear segmentation in microscope cell images: A hand-segmented dataset and comparison of algorithms. 2009 IEEE International Symposium on Biomedical Imaging: From Nano to Macro (2009). doi: 10.1109/isbi.2009.5193098

18. Smith, K. et al. CIDRE: an illumination-correction method for optical microscopy. Nature Methods 12, 404–406 (2015).

19. Naylor, P., Lae, M., Reyal, F. & Walter, T. Segmentation of Nuclei in Histopathology Images by Deep Regression of the Distance Map. IEEE Transactions on Medical Imaging 38, 448–459 (2019).

20. Kumar, N. et al. A Dataset and a Technique for Generalized Nuclear Segmentation for Computational Pathology. IEEE Trans. Med. Imaging 36, 1550–1560 (2017).

21. matterport. matterport/Mask_RCNN. GitHub Available at: https://github.com/matterport/Mask_RCNN. (Accessed: 7th October 2019)

22. Kingma, D. P. & Ba, J. Adam: A Method for Stochastic Optimization. (2014).

23. zhixuhao. zhixuhao/unet. GitHub Available at: https://github.com/zhixuhao/unet. (Accessed: 7th October 2019)

24. mirzaevinom. mirzaevinom/data_science_bowl_2018. GitHub Available at: https://github.com/mirzaevinom/data_science_bowl_2018. (Accessed: 7th October 2019)

